# Assessment of native cadmium-resistant bacteria in cacao (*Theobroma cacao* L.) - cultivated soils

**DOI:** 10.1101/2021.08.06.455168

**Authors:** Henry A. Cordoba-Novoa, Jeimmy Cáceres-Zambrano, Esperanza Torres-Rojas

## Abstract

Traces of cadmium (Cd) have been reported in some chocolate products due to soils with Cd and the high ability of cacao plants to extract, transport, and accumulate it in their tissues. An agronomic strategy to minimize the uptake of Cd by plants is the use of cadmium-resistant bacteria (Cd-RB). However, knowledge about Cd-RB associated with cacao soils is scarce. This study was aimed to isolate and characterize Cd-RB associated with cacao-cultivated soils in Colombia that may be used in the bioremediation of Cd-polluted soils. Diversity of culturable Cd-RB, qualitative functional analysis related to nitrogen, phosphorous, carbon, and Cd were performed. Thirty different Cd-RB morphotypes were isolated from soils with medium (NC, Y1, Y2) and high (Y3) Cd concentrations using culture media with 6 mg Kg^-1^ Cd. Cd-RB were identified based on morphological and molecular analyses. The most abundant morphotypes (90%) were gram-negative belong to Phylum Proteobacteria and almost half of them showed the capacity to fix nitrogen, solubilize phosphates and degrade cellulose. Unique morphotypes were isolated from Y3 soils where *Burkholderia* and *Pseudomonas* were the dominant genera indicating their capacity to resist high Cd concentrations. *P. putida* GB78, *P. aeruginosa* NB2, and *Burkholderia* sp. NB10 were the only morphotypes that grew on 18 up to 90 (GB78) and 140 mg Kg^-1^ Cd (NB2-NB10); however, GB78 showed the highest Cd bioaccumulation (5.92 mg g^-1^). This study provides novel information about culturable Cd-RB soil diversity with the potential to develop biotechnology-based strategies.

## Introduction

Cadmium (Cd) is the most prevalent toxic agent to microorganisms, plants, and human beings. It could be present in the environment naturally as a product of volcanic eruptions, weathering of bedrock, and burning of vegetation (Cullen and Maldonado 2013); or as a result of anthropogenic activities such as mining, manufacturing of plastics, paint pigments, batteries, and others (Kirkham 2006). Cd enrichment in agricultural soil is due to soil evolution (Gramlich et al. 2018) and rocks or mineral composition. Cd can be solubilized from different types of minerals such as Greenockite (CdS), Otavite (CdCO_3_), Cadmoselite (CdSe), Monteponite (CdO), and Metacinnabar Cd ([Hg, Cd]S) (Cullen and Maldonado 2013). The physical-chemical properties (e.g. texture, pH, organic matter composition, effective cation exchange capacity, total metal content), microbial community and climate conditions play also an important role (Amari et al. 2017). Additionally, agricultural practices such as the application of phosphate-based fertilizers that could reach Cd contents up to 130 mg Kg^-1^ (Siripornadulsil and Siripornadulsil 2013), crop irrigation with untreated wastewater, and contaminated raw organic material recycling for the same crop (Maddela et al. 2020) may increase Cd concentration and its mobility in soils.

The presence of Cd in plants is related to both the ability of the plant to extract, transport, and accumulate Cd in its tissues; and the availability of Cd in soil (Taki 2013). Cd has been detected in vegetables such as lettuce, swiss chard, spinach, fresh fruits, and some cereals (Quezada-Hinojosa et al. 2015). For cacao, which is considered a traditional exportable crop with a global production of 4.7 million tons (in 2018/19), where 68% comes from developing countries (ICCO, 2018; Maddela et al. 2020), Cd has also been reported (Mounicou et al. 2003; Chavez et al. 2015; Gramlich et al. 2016, 2018; Arévalo-gardini et al. 2017). The European Commission has regulated the maximum limits of permissible Cd for chocolate and other derived products depending on the percentage of raw cacao present in the final product (Commission regulation (EU) No. 488/2014 2014) where the levels range from 0.1 – 0.8 mg Kg^-1^ Cd dry matter (Meter et al. 2019). Concentrations of Cd in cacao-based chocolate and raw cacao beans above these critical levels have been reported in some South and Centro American countries (Chavez et al. 2015; Bertoldi et al. 2016). This situation represents a major concern because of the impact on food chain that compromises human health (Clemens et al. 2013), food safety, and the cacao economy (Maddela et al. 2020).

It has been reported different agronomic strategies to reduce Cd uptake by plants that include the management of pH, organic matter content, mineral nutrition, and replacement of contaminated fertilizers (Rai et al. 2019). The reduction of Cd levels can also be achieved through the application of biological amendments. The use of microorganisms with the ability to adsorb, bioaccumulate and biotransform Cd is an attractive approach to carry out bioremediation processes of contaminated soils (Ashraf et al. 2017). Soil bacteria have several resistance mechanisms to overcome toxic levels of Cd such as biosorption, intracellular sequestration, extracellular binding, complexation, and removal by efflux systems. A reduction of Cd uptake by plants using Cd resistant bacteria (Cd-RB) and Cd resistant plant growth-promoting rhizobacteria (PGPR) has been reported in wheat and maize (Ahmad et al. 2014), rice (Treesubsuntorn et al. 2017), mustard (Sinha and Mukherjee 2008), tomato (Madhaiyan et al. 2007), pea (Safronova et al. 2006), and soybean (Guo and Chi 2014). Among Cd-RB with the potential to reduce the availability of this heavy metal at the interchangeable soil phase, *Pseudomonas aeruginosa* (Chakraborty and Das 2014), *Pseudomonas putida* (Li et al. 2014), *Serratia* spp. (Sarma et al. 2016), *Burkholderia* spp. (Jin et al. 2013), *Halomonas* spp. (Asksonthong et al. 2016) and *Enterobacter* spp. (Chen et al. 2016) have been commonly reported.

In Colombia, where the spatial distribution and concentration of Cd in cacao-cultivated soils has been mainly associated with a geogenic origin (Rodríguez Albarrcín et al. 2019; Bravo and Benavides-Erazo 2020), Cd-RB belonging to genera *Enterobacter* sp., *Burkholderia* sp., *Pseudomonas* sp., and *Methylobacterium* sp. from cacao-cultivated soils with Cd have been isolated (Bravo et al. 2018). However, the knowledge of the taxonomic diversity, community structure, and functional activity of Cd-RB associated with cacao crop are still scarce. The purpose of this study was to isolate and characterize culturable Cd-RB associated with cacao-cultivated soils with Cd, to contribute to the knowledge of this bacterial community that may be used in integral soil management programs.

## Materials and methods

### Study area and sampling

Soil samples were obtained from two municipalities of Cundinamarca, Colombia: Nilo (NC) 4°18’25’’N - 74°37’12’O, and Yacopí (Y) 5°27’35’’N - 74°20’18’O (Rodríguez Albarrcín et al. 2019). These regions have an average temperature of 26.5 °C and 21°C with annual precipitation of 1292 and 1500 mm, respectively. Three cacao-producing farms were sampled in Y and one in NC (Fig 1). In each sampling site, three producing trees were randomly selected, and four soil sub-samples (0.5 Kg each) were collected at 0.3 m deep with 0.5 m from the trunk base. Sub-samples were mixed to get a composite sample per farm. These samples were used for the determination of physical and chemical properties, and taxonomic classification (Supplemental Table 1; Table 1).

**Table 1.**
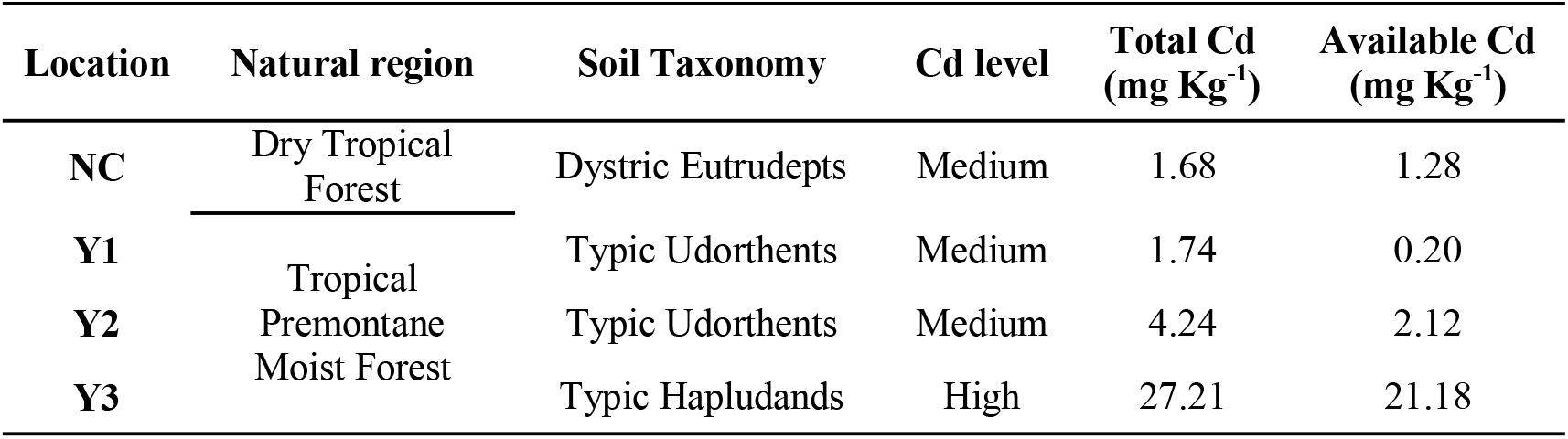
Location, soil taxonomy, and total and potentially available Cd concentrations in sampling sites in Nilo (NC) and Yacopí (Y). Medium (>1.0 to 5.0 mg Kg^-1^) and high (>5.0 mg Kg^-1^) total Cd level

**Fig. 1.**
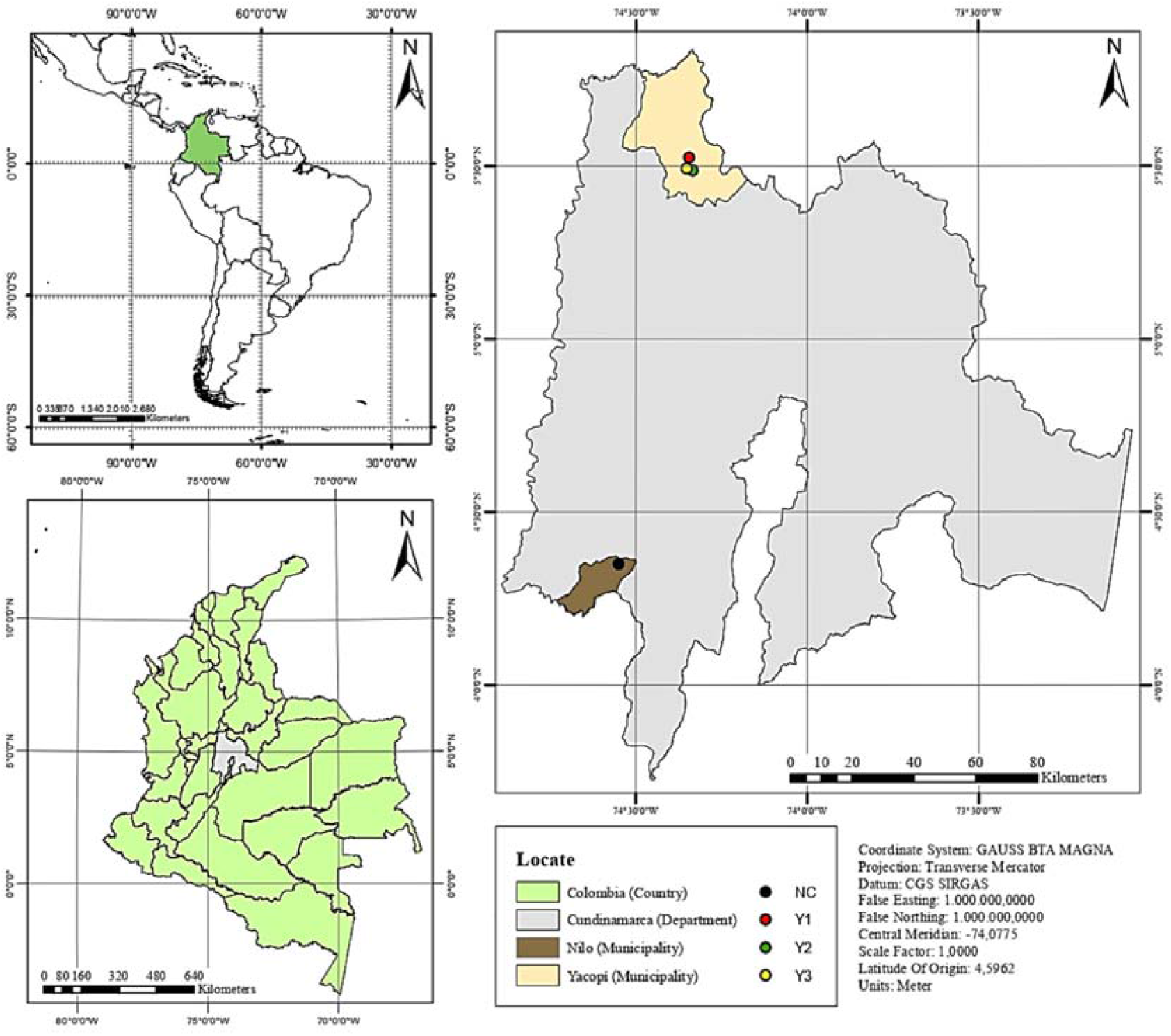
Location of sampling sites. Each dot represents a cacao farm where previously soil cadmium levels have been documented

### Soil cadmium determination

Composite samples were used to quantify the total Cd and potentially available Cd. Total Cd was determined by digestion of 1 g of soil sample with 7 mL of HNO_3_ 65% (v/v) and 1 mL of HCl 37% (v/v). Samples and acids were mixed using a single reaction chamber UltraWAVE (ECR; Milestone, Shelton, CT), the temperature rose from 20 °C to 220 °C in 20 min and was maintained for 10 min for a total digestion time of 30 min. After digestion, 10 mL of deionized water was added, and samples were filtered using 0.2 µm x 47 mm sterile Swinnex filters (Merck-Millipore, Darmstadt, Germany). Potentially available Cd was quantitatively determined with DTPA extraction solution [0.005 M diethylenetriaminepentaacetic acid (DTPA), 0.01 M calcium chloride dihydrate (CaCl_2_ 2H_2_O) and 0.1 M triethanolamine (TEA)] in a 1:2 proportion (w/v).

Determinations were performed using an Agilent 700 series Inductively Coupled Plasma Optical Emission Spectroscopy (ICP-OES) spectrometer (Agilent Technologies, Inc., CA, USA) with five milliliters of the digested sample and a running time of 90 s per sample. The ICP Expert II software was used to analyze raw ICP data. A standard curve was created using CdCl_2_ aliquots (Sigma-Aldrich Corp., 99% purity w/w, CA) in eight concentrations ranging from 0 to 500 mg L^-1^; Supplemental Fig. 1). Additionally, aliquots of the reference material WEPAL-ISE-997 (Sandy soil with 0.400 mg Kg^-1^ of Cd^2+^, obtained from the Wageningen evaluating program for analytical laboratories, Wageningen Agricultural University, the Netherlands) were included (Bravo et al. 2018).

In Colombia, neither reference values for soil Cd are available for agricultural production. In non-polluted soils, Cd concentrations range from 0.01 to 1.1 mg Kg^-1^ with an average of 0.41 mg Kg^-1^ (Kabata-Pendias and Pendias 2011). According to this, in this study Cd levels in the four locations were categorized as medium (>1.1 to 5.0 mg Kg^-1^) and high (>5.0 mg Kg^-1^; Table 1).

### Culturable Cd-RB isolation, morphological characterization, and molecular identification

Cd-RB were isolated by plating out serial dilutions of soil as described previously (Avellaneda-Torres et al. 2020) with slight modifications. 50 g of sifted soil (0.2 mm) were suspended in 450 mL of NaCl 0.85%, mixed for 10 min and, 1 mL of the dilution was transferred to a new assay tube with 9 mL of NaCl and mixed with vortex for 1 min. The process was made up to 10^−7^ dilution and 100 µl of the dilutions 10^−3^, 10^−5^ and, 10^−7^ were plated out on Mergeay agar (Mergeay 1995) pH 7 with 6 mg Kg^-1^ of Cd from CdCl_2_ and incubating at 37 °C for 3 days. Following incubation, the colony-forming unit (CFU) was calculated per gram of dry soil as an indirect measure to determine total abundance by location (Bressan et al. 2015). Morphotypes with differential morphological characteristics were subcultured in Mergeay agar with 6 mg Kg^-1^ of CdCl_2_ and were preserved in glycerol stocks at - 70 °C.

Micro and macro morphological characterization were performed to each isolated assessing color using the Pantone scale, shape, elevation, surface appearance, and colony consistency. Also, Gram-stain and endospore stain with malachite green dye in contrast with safranin were performed. Molecular identification was made by extracting DNA according to Chen y Kuo (Chen and Kuo 1993) and partial amplification of the 16S rRNA gene with the universal primers 27F and 1492R as previously described (Avellaneda-Torres et al. 2020). All sequences were submitted to the GenBank (NCBI).

### Analysis of structural diversity and phylogeny of culturable Cd-RB

Cd-RB structural diversity was assessed by (i) richness and abundance; richness was considered as the number of present genera per sample and the abundance as the number of isolated morphotypes for each location (NC, Y1, Y2, Y3) (Lyngwi et al. 2013; dos Passos et al. 2014) and (2) phylogenetic analysis of the 16 rRNA genes from isolates. The phylogenetic tree was inferred using the *Muscle* multiple alignment method and *Neighbor-joining* algorithm (Saitou and Nei 1987). The evolutionary distances were computed using the p-distance method and nodes are supported in the bootstrap method using 1000 pseudoreplicates (Felsenstein 1985). Evolutionary analyses were conducted in MEGA7 (Kumar et al. 2016).

### Functional analysis of isolated Cd-BR morphotypes associated with C, P, and N

The cellulolytic potential of isolated morphotypes was assessed in LB agar with carboxymethyl cellulose CMC, as the unique carbon source, and staining with Congo red (Avellaneda-Torres et al. 2014). Phosphate solubilization activity was assessed in SMRS agar (Paul and Sundara 1971), and the solubilization factor (SF) was calculated using bromocresol purple. Hydrolysis and solubilization factors establish the ratio of degradation or solubilization diameter and colony growth diameter (Paul and Rao 1971). Finally, atmospheric nitrogen fixation activity was observed on Rennie agar (Rennie 1981) with bromothymol blue as an indicator, through growth and turning color of this medium that lacks N. Three replicates per morphotype were used for each functional group.

### Effect of Cd on bacterial growth, Cd bioaccumulation, and Minimum Inhibitory Concentration

Morphotypes previously isolated on Mergeay agar with 6 mg Kg^-1^ of Cd were inoculated in LB agar with 12 and 18 mg Kg^-1^ of Cd. Bacterial growth curves were analyzed for bacteria resistant to 18 mg Kg^-1^ of Cd. LB broth with (18 mg Kg^-1^) and without Cd was inoculated with 0.1% (v/v) of bacterial cultures of 18 h (Ivanova et al. 2002; Cristani et al. 2012). Bacteria were incubated at 37 °C with shaking at 200 rpm. With three replicates for each morphotype, optic density at 600 nm (OD600) was determined every 2 h during 36 h using a NanoDrop™ ONE spectrophotometer (ThermoFisher Scientific, USA).

Cd bioaccumulation test was made twice at different moments. The bacteria were inoculated into LB broth containing 18 mg Kg^-1^ of Cd and incubated at 37 °C with shaking (200 rpm) with three replicates per bacteria. The bacterial cells were harvested at 18 h after inoculation by centrifugation at 5000 rpm for 15 min and then washed with sterile deionized water to remove free heavy metal ions. The supernatants were sterilized by filtration with a 0.22 µm membrane filter (MILLIEX^®^GP Millipore Express^®^) and cell pellets were dried at 100 °C until constant weight. Media without adding Cd or inoculating bacteria were used as controls. The concentration of Cd in supernatants was measured by Atomic Absorption Spectrophotometry (AAS; Perkin Elmer AAnalyst 300, USA) and the metal bioaccumulation of bacteria was calculated as follows (Micheletti et al. 2008):

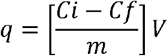

where *q* (mg g^-1^) is the Cd bioaccumulation of bacterial cells, *Ci* is the initial concentration of heavy metal used (mg L^-1^), *Cf* is the final concentration of heavy metal (mg L^-1^), *m* is the dried weight of cell pellet (g) and *V* is the volume of the liquid medium (0.05 L in the experiments).

The bacteria resistant to 18 mg Kg^-1^ of Cd were grown in 15 mL conical bottom tubes with 7 mL of LB broth supplemented with 18, 24, 30, 40 up to 200 mg Kg^-1^ of Cd in intervals of 10 mg Kg^-1^ of Cd. 20 µl of bacterial culture of 18 h were inoculated and incubated at 37 °C with constant agitation at 200 rpm for 18 h. Following incubation, turbidity was observed. The minimum Cd concentration at which bacterial isolate did not show growth was considered as its Minimum inhibitory concentration (MIC).

### Data analysis

Data from hydrolysis and solubilization factors and Cd bioaccumulation were analyzed using a completely randomized design (CRD) with a one-way analysis of variance (ANOVA) in SAS v9.4 statistical software (SAS Institute Inc., USA). Shapiro-Wilk normality test and Bartlett homogeneity of variance tests were made. Media were compared using Tukey’s test (p*<* 0.05).

## Results

### Culturable of Cd-RB isolation, morphological characterization, and molecular identification

A total of 30 Cd-RB morphotypes were isolated from the different soil samples in culture media with 6 mg kg^-1^ Cd. One isolate of each morphotype was morphologically characterized and identified. The number of CFU of Cd-RB was less than 10^6^ CFU g^-1^ dry soil in all locations without significant differences (Supplemental Fig. 2). The micro and macroscopic characterization are shown in Supplemental Table 2, and the Gram analysis showed that 90% of the isolated (n=27) corresponded to Gram-negative bacteria and the remaining 10% (n=3) to Gram-positive (Table 2). Based on the BLAST search results of the 16S rRNA gene, it was identified eight genera: *Burkholderia, Pseudomonas, Enterobacter, Serratia, Bacillus, Halomonas, Herbaspirillum*, and *Rhodococcus* (Supplemental Table 3).

**Table 2.** Cellulose hydrolysis and phosphate solubilization factors, and free nitrogen-fixing activity of Cd-RB. Letters in the same columns represent significant differences (*p*<0.05) between strains according to Tukey’s multiple comparison test. + and – indicate the presence or absence of growth/fixation on Rennie agar at 3 days post-inoculation (dpi)

| No. | ID | NCBI ID | Loc. | Morphotype | Growth at |  | Cellulose hydrolysis factor | P solubilization factor | N fixation |
| --- | --- | --- | --- | --- | --- | --- | --- | --- | --- |
|  |  |  |  |  | 12 mg Kg<sup>-1</sup> | 18 mg Kg<sup>-1</sup> |  |  |  |
| 1 | NB2 | MN719036.1 | NC | *P. aeruginosa* | + | + | - | - | - |
| 2 | NB10 | MN719037.1 | NC | *Burkholderia* sp. | + | + | 2.04 ± 0.28 abcd | 3.31 ± 0.55 abc | + |
| 3 | NB40 | MN719038.1 | NC | *Halomonas* sp. | + | - | - | - | - |
| 4 | NB58 | MN719039.1 | NC | *Enterobacter* sp. | + | - | - | - | - |
| 5 | NB59 | MN719040.1 | NC | *Serratia* sp. | + | - | - | - | - |
| 6 | NB61 | MN719041.1 | NC | *Serratia* sp. | + | - | - | < 1.00 | - |
| 7 | NB80 | MN719042.1 | NC | *Serratia* sp. | + | - | 1.92 ± 0.23 bcd | < 1.00 | - |
| 8 | YB11 | MN719043.1 | Y1 | *Enterobacter* sp. | + | - | 3.25 ± 0.28 a | 3.83 ± 0.48 ab | + |
| 9 | YB30 | MN719044.1 | Y1 | *S. marcescens* | + | - | - | 4.39 ± 0.49 a | + |
| 10 | YB42 | MN719045.1 | Y1 | *S. marcescens* | + | - | - | - | - |
| 11 | YB43 | MN719046.1 | Y1 | *Serratia* sp. | + | - | 0.99 ± 0.01 d | < 1.00 | + |
| 12 | YB5 | MN719047.1 | Y2 | *Enterobacter* sp. | + | - | 1.53 ± 0.30 cd | 1.56 ± 0.21 c | + |
| 13 | YB13 | MN719048.1 | Y2 | *Pseudomonas* sp. | + | - | - | - | - |
| 14 | YB22 | MN719049.1 | Y2 | *Bacillus flexus* | - | - | - | - | - |
| 15 | YB48 | MN719050.1 | Y2 | *Enterobacter* sp. | + | - | 2.00 ± 0.08 bcd | 2.02 ± 0.18 bc | + |
| 16 | GB13 | MN719051.1 | Y3 | *Burkholderia* sp. | + | - | - | < 1.00 | + |
| 17 | GB16 | MN719051.1 | Y3 | *Burkholderia* sp. | + | - | 1.04 ± 0.13 d | < 1.00 | + |
| 18 | GB17 | MN719053.1 | Y3 | *Rhodococcus* sp. | + | - | 1.01 ± 0.04 d | < 1.00 | + |
| 19 | GB18 | MN719054.1 | Y3 | *Bacillus* sp. | + | - | - | - | - |
| 20 | GB58 | MN719055.1 | Y3 | *Burkholderia* sp. | + | - | 1.07 ± 0.11 d | - | - |
| 21 | GB66 | MN719056.1 | Y3 | *Burkholderia* sp. | + | - | 1.20 ± 0.02 d | < 1.00 | + |
| 22 | GB67 | MN719057.1 | Y3 | *Burkholderia* sp. | + | - | - | - | - |
| 23 | GB68 | MN719058.1 | Y3 | *Burkholderia* sp. | + | - | 2.73 ± 0.58 abc | 3.46 ± 0.46 abc | + |
| 24 | GB71 | MN719059.1 | Y3 | *Pseudomonas* sp. | + | - | - | < 1.00 | + |
| 25 | GB73 | MN719060.1 | Y3 | *Pseudomonas* sp. | + | + | - | - | - |
| 26 | GB78 | MN719061.1 | Y3 | *P. putida* | + | + | 1.18 ± 0.01 d | < 1.00 | + |
| 27 | GB82 | MN719062.1 | Y3 | *Burkholderia* sp. | + | - | - | - | - |
| 28 | GB86 | MN719063.1 | Y3 | *Pseudomonas* sp. | + | - | 3.12 ± 0.43 ab | < 1.00 | + |
| 29 | GB88 | MN719064.1 | Y3 | *Pseudomonas* sp. | + | - | - | - | - |
| 30 | GB90 | MN719065.1 | Y3 | *Herbaspirillum* sp. | + | - | 2.77 ± 0.21 abc | - | + |

**Fig. 2.**
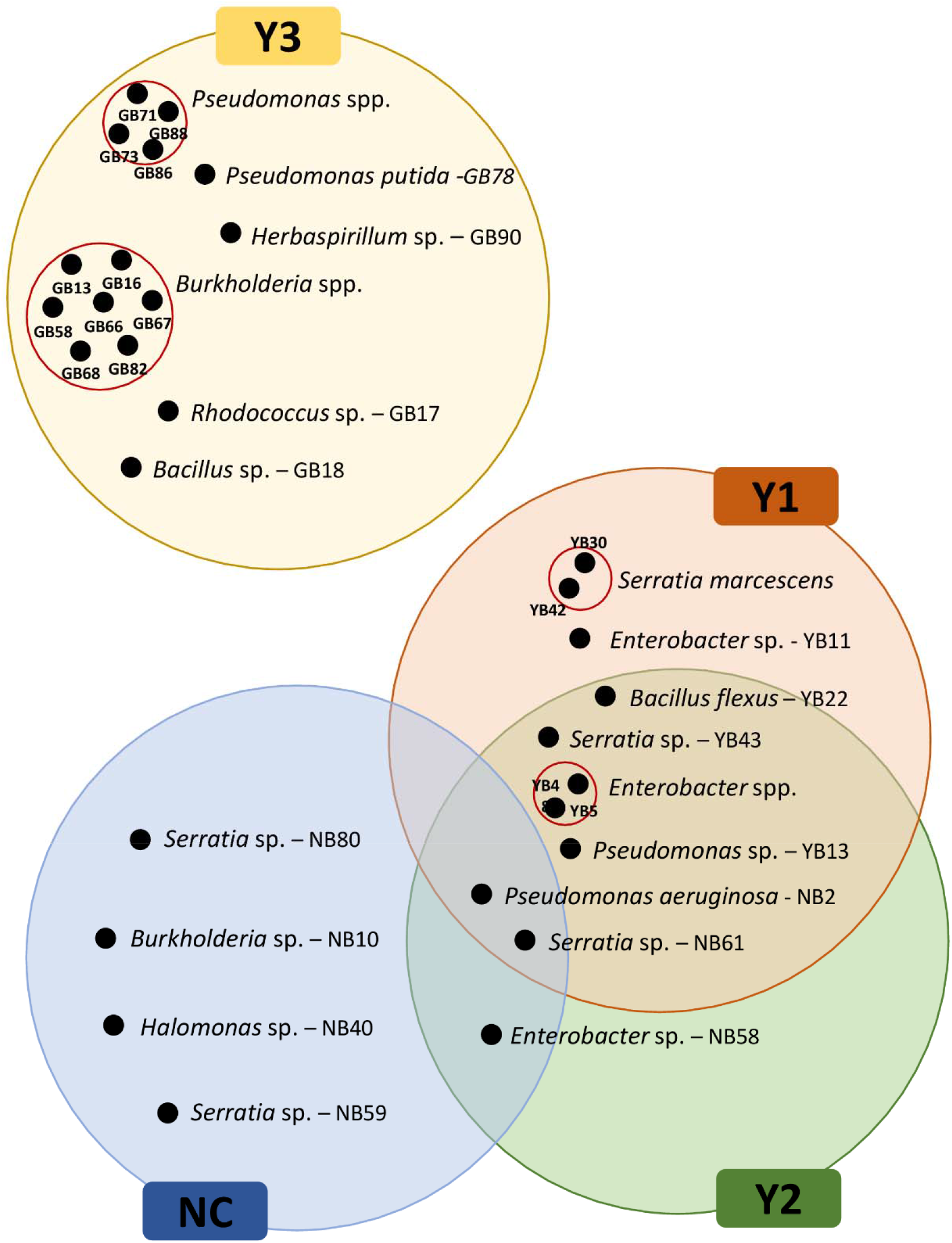
Venn diagram of the culturable Cd-RB morphotypes isolated from locations studied. Each gray circle represents the location, referred to with a label. Black circles represent the isolated morphotypes, the different morphotypes belonging to the same genera are represented by the red contour circle. The morphotypes were isolated from cacao cultivated soils with medium (>1.0 to 5.0 mg Kg^-1^; CN, Y1 and Y2) and high (>5.0 mg Kg^-1^; Y3) total Cd levels

### Analysis of structural diversity and phylogeny of culturable Cd-RB

It also was found a greater abundance, unique morphotypes, and dominance of isolated Cd-RB belonging to the genera *Burkholderia* and *Pseudomonas* in soil with the highest natural concentration of Cd (location Y3; Fig. 2). Additionally, Y1 and Y2 shown the greatest similarity according to the isolated morphotypes (Fig. 2).

The phylogenetic analysis showed that 90% (n=27) of the isolated bacteria belong to the phylum Proteobacteria, 6.7% to Firmicutes (n=2), and 3.3% to Actinobacteria (n=1) (Fig. 3; Supplemental Table 3). In the phylogenetic tree, *Rhodococcus* sp. GB17, *Bacillus flexus* YB22, and *Bacillus* sp. GB18 were used as an outgroup due to their relationship with the other isolated morphotypes. Within the phylum proteobacteria, two classes stand out, Gamma and Beta Proteobacteria. Gammaproteobacteria were represented by the genera *Serratia* (6 morphotypes), *Enterobacter* (4 morphotypes), *Pseudomonas* (7 morphotypes), and *Halomonas* (1 morphotype) belonging to the families Enterobacteriaceae (Enterobacteriales), Pseudomonadaceae (Pseudomonadales), and Halomonadaceae (Oceanospirillales), respectively. Beta Proteobacteria had the genera *Burkholderia*, which presented the highest number of morphotypes found (8), and *Herbaspirillum* (1); belonging to the family Burkholderiaceae and Oxalobacteraceae, respectively, and both to order Burkholderiales. The genera *Bacillus* (2 morphotypes) and *Rhodococcus* (1) were the least represented, belonging to the Bacillaceae (Bacillales) and Nocardiaceae (Actinomycetales) class, respectively.

**Fig. 3.**
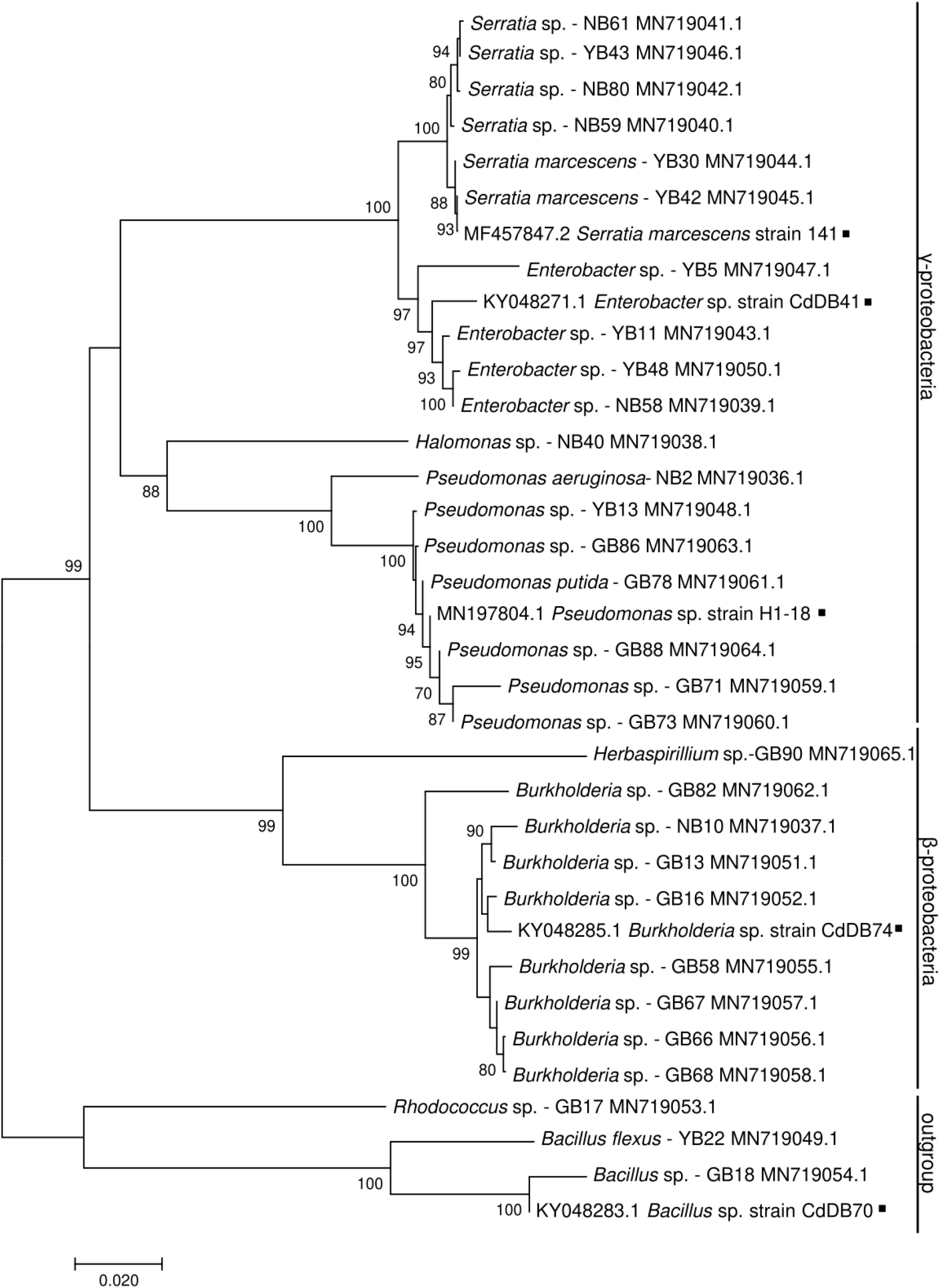
Phylogenetic tree of strains isolated in Mergeay agar with 6 mg kg^-1^ of Cd from cacao culture soil. The evolutionary history was inferred using the *neighbor-joining* method. The bootstrap test was inferred with 1000 replicates and the analysis was performed with 36 sequences. Positions containing gaps and missing data were eliminated. The evolutionary distances were computed with the p-distance method. Evolutionary analyses were performed in MEGA 7. Sequences indicated with black squares were used as a template

### Functional analysis of isolated morphotypes associated with C, P, and N

Some isolated bacteria presented functional activities related to C, P, and N in addition to their Cd resistance. The 46.7% (n=14) of the isolated morphotypes showed the ability to degrade cellulose, highlighting strains such as *Enterobacter* sp. YB11, *Pseudomonas* sp. GB86, *Herbaspirillun* sp. GB90 and *Burkholderia* sp. GB68 with hydrolysis factors above 2.5 (Table 2). Regarding P, a higher percentage of morphotypes were able to solubilize phosphates (53%, n=16) compared with other functional groups analyzed. Bacteria from genera *Enterobacter, Burkholderia*, and *Serratia* showed high solubilization factors ranging from 1.56 to 4.39 (Table 2). Finally, atmospheric nitrogen fixation ability was observed in half of the isolated morphotypes.

### Effect of Cd on bacterial growth, Cd bioaccumulation, and Minimum Inhibitory Concentration

Among the 30 isolated morphotypes only *Pseudomonas* sp. GB73, *Burkholderia* sp. NB10, *P. aeruginosa* NB2, *P. putida* GB78 grew on media with 18 mg Kg^-1^ Cd, and the last three were considered for further analysis. The effect of Cd on the growth curve varied among the assessed strains. In presence of 18 mg Kg^-1^, *P. aeruginosa* NB2 showed an average reduction of more than one unit of OD_600_ during the log phase from 14 h after inoculation compared to control without Cd (Fig. 4A). In contrast, *P. putida* GB78 did not show any difference in OD_600_ with and without Cd during all growth phases (Fig. 4B). *Burkholderia* sp. NB10 had a slight reduction of OD_600_ with Cd from 2 to 20 h post-inoculation, however, from 20 h the OD_600_ of bacteria grown with Cd reached similar magnitudes to culture without Cd.

**Fig. 4.**
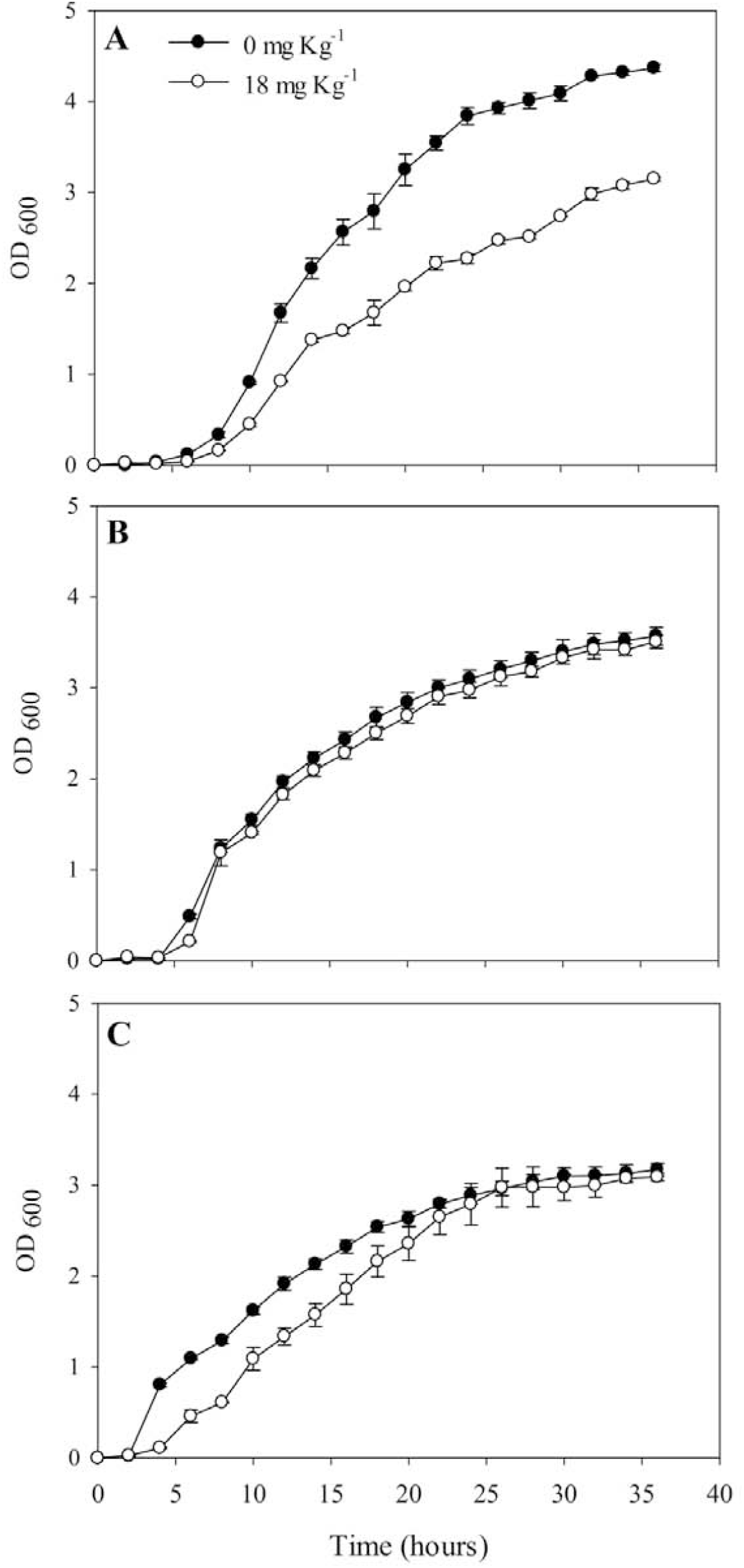
Growth of A) *P. aeruginosa* NB2, B) *P. putida* GB78 and C) *Burkholderia* sp. NB10 in the presence and absence of 18 mg Kg^-18^ of Cd. The bars represent the standard error (n=6)

Regarding Cd bioaccumulation (*q*), *P. putida* GB78 accumulated 5.92 mg g^-1^, which was significantly higher than *P. aeruginosa* NB2 and *Burkholderia* sp. NB10 accumulation (around 1 mg g^-1^; Fig. 5). Finally, it was found that *Burkholderia* sp. NB10 and *P. aeruginosa* NB2 showed a MIC of 140 mg Kg^-1^, which is 1.5 times higher than the MIC recorded for *P. putida* GB78 (90 mg Kg^-1^).

**Fig. 5.**
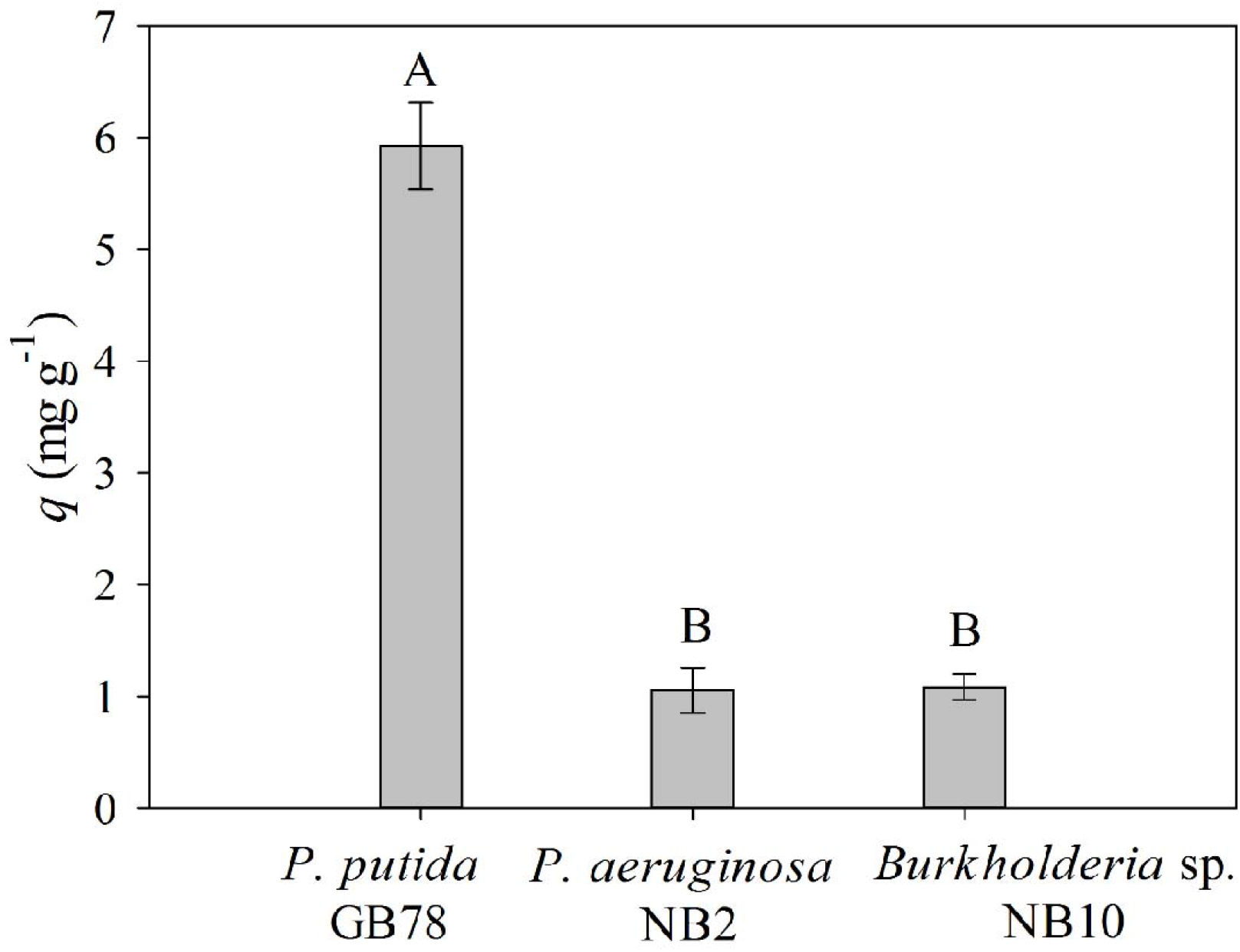
Bioaccumulation of Cd in bacteria at 18 h of growth on media with 18 mg Kg^-1^ of Cd. The bars represent the typical error (n=3). Means with different letters indicate significant differences by Tukey’s test (*p*<0.05)

## Discussion

### Culturable of Cd-RB isolation, morphological characterization, and molecular identification

The Log CFU g^-1^ of dry soil is an indirect measure of the abundance of microorganisms present in soil and for agricultural soils, values between 10^8^ – 10^10^ cells per gram have been reported (Marteinsson et al. 2015). In this study, the bacterial abundances for all studied locations were considered low (below 10^6^), which could be attributed to the presence of Cd as a selection factor in the isolation culture media with 6 mg Kg^-1^ Cd (Torsvik et al. 2002; Itävaara et al. 2016). Although it has been found that heavy metals affect adversely the composition of the microbial community (Etesami 2018), in this study, it was selected cacao-cultivated soil and culture media with a high concentration of Cd to isolated bacteria, due to these conditions would allow a better selection of morphotypes with an enhanced capacity to survive under Cd stress condition *in vitro*.

The morphological characterization showed that 90% of the isolated morphotypes corresponded to Gram-negative bacteria. Gram-negative and Gram-positive bacteria respond differently to the presence of heavy metals (Bruins et al. 2000; Limcharoensuk et al. 2015). Bruins et al. (Bruins et al. 2000) assert that some Gram-negative bacteria (genus *Pseudomonas*) can tolerate 5 to 30 times more Cd than Gram-positive bacteria (*Staphylococcus aureus, S. faecium, Bacillus subtilis*). In this regard, Wu et al. (Wu et al. 2009) reported that *Azotobacter chroococum*, a Gram-negative bacteria, had a greater capacity for binding to Cd than *Bacillus megaterium*, a Gram-positive one. This differential response can be explained due to the composition of the cell wall and the resistance mechanisms used.

Gram-positive bacteria have a thick layer of peptidoglycan (20-80 nm) in which they fix acids, proteins, and polysaccharides. This layer forms a barrier to avoid the entry of heavy metals into the cell, prevailing passive resistance mechanisms such as adsorption (Bruins et al. 2000; Mounaouer et al. 2014). In contrast, Gram-negative bacteria have thinner walls (5-10 nm) that allow the influx and efflux of heavy elements and force the bacteria to develop active mechanisms or flow systems and to have a higher protein synthesis in presence of Cd (Bruins et al. 2000). Nevertheless, both types of bacteria can present passive and active resistance mechanisms to Cd and other heavy metals at the same time (Sharma and Archana 2016; Etesami 2018). This indicates that resistance depends also on the other factors such as type of strain, type, and concentration of the heavy metals, physicochemical properties of soil, and environmental conditions (Sharma and Archana 2016). In this regard, Lima et al. (Lima e Silva et al. 2012) observed a greater presence of Gram-positive Cr resistant bacteria, while Hg and Ag resistant bacteria were mostly Gram-negative. This demonstrates that the pollutant also determines the type of bacteria that survive in contaminated environments.

*Burkholderia, Pseudomonas, Enterobacter*, and *Serratia* were the most frequent isolated genera. These genera have been reported in the management of cultivated soils. Strains of *Burkholderia* spp. have the potential to promote plant growth, biocontrol pathogens, and biodegrade toxic molecules (Castanheira et al. 2016). *Enterobacter* spp. can improve bioremediation processes of soils contaminated with heavy metals (Qiu et al. 2014; Marteinsson et al. 2015) and within the genus *Pseudomonas*, growth-promoting strains have also been reported with the ability to mitigate stress in plants caused by the presence of heavy metals such as Cd and Pb, and metals such as Zn that are toxic at high levels (Lin et al. 2016; Rojjanateeranaj et al. 2017). *Serratia* sp. has been reported as a bacteria that can confer tolerance to heavy metal stress in cereals (Ahmad et al. 2014). Additionally, genera found in less frequency such as *Halomonas* and *Herbaspirillum*, have been involved in bioremediation processes of Cr and Sr in symbiosis with algae, and in free life (Achal et al. 2012; Murugavelh and Mohanty 2012; Gupta and Diwan 2017), with the potential to fix nitrogen, and chelate heavy metals such As, Pb, and Cu (Govarthanan et al. 2014; Li et al. 2018). The bacteria isolated in this study, constitute a potential resource of native microorganisms to manage Cd in cacao cultivated soils. However, further studies to characterize and evaluate the performance of bacteria in the field and their interaction with plants, are needed.

### Analysis of structural diversity and phylogeny of culturable Cd-RB

The structure of microbial communities can be affected by the presence of heavy elements. In this study, results showed that there was not a relationship between the richness of isolated morphotypes and the level of Cd present in the soil. However, the presence of unique morphotypes and the dominance of *Burkholderia* and *Pseudomonas* genera (Fig. 2), show that the presence of Cd in soils may alter the uniformity of the community of Cd-RB present in the studied soil. It has been reported that more sensitive microorganisms to heavy metals are suppressed by the toxic element and replaced by more resistant with adaptive characteristics, occupying their ecological niche (Azarbad et al. 2015). The presence of unique morphotypes isolated from Y3 soils agrees with the results reported by Tipayno et al. (Tipayno et al. 2018) who studied sites with different levels of Cd and other heavy metals (As and Pb) and found that the site with the highest concentration of Cd also presented the highest number of unique morphotypes.

Most of the isolated morphotypes (90%) in this study corresponded to the Proteobacteria phylum. This phylum represents the largest and most diverse group of prokaryotes and contemplates the vast majority of Gram-negative bacteria studied to date (Spain et al. 2009). Also, it is usually reported as the most frequent in studies related to heavy metals, so strains belonging to several genera have been isolated, characterized, and reported by their resistance (Xiao et al. 2019). Additionally, Chen et al. (Chen et al. 2018) highlight that the Proteobacteria phylum together with Bacteroidetes and Firmicutes harbor the largest set of heavy metal resistance genes.

### Functional analysis of isolated Cd-RB morphotypes associated with C, P, and N

Cellulolytic potential, free nitrogen fixation, and phosphate solubilization were determined in all isolated morphotypes using selective media to promote an integrated crop cacao management. The use of these bacteria could enhance nutrient cycling, decrease the use of chemical fertilizers, and increase the environmental sustainability of the crop. In agroforestry production systems such as cacao, some interactions and processes favor nutrient cycling, as a result of the decomposition of organic matter from crop residues and pruning such as litter in the crop, stems, among others (Aikpokpodion 2010). The decomposition of cellulose increases the levels of organic matter, and carbon can be used as a source of energy for edaphic processes and plant physiology (Gupta et al. 2012). Organic matter is a fraction to which heavy metals such as Cd can bind and be adsorbed by chelates or other substances. Therefore, when levels of organic matter increase in soil, the bioavailability of Cd for plants can be reduced (Gadd 2000).

The solubilization of phosphates and the promotion of plant growth have been reported for the genus *Enterobacter* isolated from soil (Beltrán 2014). The main cellular mechanism through which bacteria solubilize phosphates is the production of organic acids such as gluconic acid, 2-ketogluconic acid, lactic acid, isovaleric acid, isobutyric acid, and acetic acid (Beltrán 2014). These molecules have a negative charge and therefore can form complexes with metal cations in solution and be involved in the chelation of Cd (Yang et al. 2018). The bacteria involved in the atmospheric nitrogen fixation (ANF-B) have also played an important role in the remediation of Cd polluted soils (Chen et al. 2020). Ivishina et al. (Ivshina et al. 2014) showed that ANF-B can secrete extracellular polymeric substances (EPS) and various organic acids and amino acids that may alter the Cd availability in the soil by adsorbing and trapping metals due to the presence of many anionic functional groups.

The management of Cd-polluted agricultural soils is a complex issue where the Cd-RB could have an important function not only to decrease the Cd availability but also to improve soil fertility. The presence of activity associated with the cycling of C, P, and N in Cd-RB is an advantage within holistic crop soil management strategies. In this study, 11 from 30 isolated morphotypes showed the ability to degrade cellulose, solubilize phosphate and, fix atmospheric nitrogen, simultaneously. It would be valuable additional analyses that quantify the activity of involved enzymes and determine the effect of bacteria on plant growth and physiology.

### Effect of Cd on bacterial growth, Cd bioaccumulation, and Minimum Inhibitory Concentration

The effect of Cd on bacterial growth varies depending on bacterial strain. Heavy metals affect bacterial growth by causing multiple metabolic alterations such as DNA damage, decreasing protein synthesis, disruption of respiration and cell division, and amino acid biosynthesis (Chen et al. 2016). These effects explain the growth decrease of *P. aeruginosa* NB2 at 18 mg Kg^-1^ of Cd along with the growth phases (Fig. 4A). In *Burkholderia* sp. NB10, a reduction of the growth was also detected in the lag and log phases (Fig. 4C); however, the cells were able to adapt and resume growth at the stationary phase. It can be explained by the repair of cadmium-mediated cellular damage and adjustment of the cell physiology to limit the distribution of toxic ions in the cell (Mohamed and Abo-Amer, 2006).

Despite Cd effects, bacteria have different Cd resistance strategies dependent and independent on intracellular accumulation (Ashraf et al. 2017; Etesami 2018). Mechanisms of resistance include metal efflux systems, transportation, precipitation, transformation, and sequestration by metallothionein proteins and thiol compounds such as glutathione (Nies 2003). The results showed that resistance mechanisms of *P. putida* GB78 are efficient enough to resist 18 mg Kg^-1^ of Cd without affecting its growth. These results can also be explained based on the difference between the efficiency of resistance mechanisms of strains. Chen et al. (Chen et al. 2014) suggest that bacteria can activate resistance mechanisms such as metal efflux bombs at particular moments which results in a depletion of intracellular Cd content.

The evaluation of Cd bioaccumulation (*q*) is a measure of the potential of microorganisms to be used in bioremediation processes and management of toxic concentrations of heavy metal in agricultural soils (Cánovas et al. 2003). A higher *q* detected in *P. putida* GB78 suggests that this strain favors intracellular accumulation processes (Fig. 5). In contrast, *Burkholderia* sp. NB10 and *P. aeruginosa* NB2 showed a lower *q*, which may be attributed to greater efficiency in extracellular Cd outflow mechanisms.

The MIC is an indication of the resistance capacity that a microorganism has to grow in the presence of Cd and highlights its potential for applications at different heavy metal concentrations in soils (Xu et al. 2012). Results showed a higher resistance in *Burkholderia* sp. NB10 and *P. aeruginosa* NB2 (140 mg Kg^-1^) compared to *P. putida* GB78 (90 mg Kg^-1^). Different *Pseudomonas* and *Burkholderia* species have been widely reported for their resistance to Cd, other heavy metals such as Cu and Pb, and high concentrations of Zn (Naik and Dubey 2011; Jin et al. 2013). It has been reported MICs from 100 to 190 mg Kg^-1^ for *P. putida* (Lee et al. 2001; Leedjärv et al. 2008), up 900 mg Kg^-1^ for *P. aeruginosa* (Ghaima et al. 2017), and 200 mg Kg^-1^ for *Burkholderia* sp. (Jin et al. 2013). These values show the wide variation that exists in the level of resistance in bacteria. Overall, the bacterial growth and the MIC results suggest that 18 mg Kg^-1^ of Cd is not a limiting concentration for *P. putida* GB78. However, it can resist up to 90 mg Kg^-1^ of Cd by favoring intracellular bioaccumulation. On the contrary, *Burkholderia* sp. NB10 and *P. aeruginosa* NB2, with a MIC of 140 mg Kg^-1^ of Cd, showed growth rate reductions that may be an acclimatization strategy to maintain viability for a longer time and activate resistance mechanisms such as efflux systems (lower *q*).

## Conclusions

Cacao-cultivated soils harbor a diversity of bacteria that exhibit resistance to Cd and functional activity associated with the cycling of C, P, and N. The structural diversity of culturable Cd-RB bacterial community is affected by the presence of Cd in soil samples where high levels of Cd enhance the selection of unique and dominant morphotypes such as *Burkholderia* spp. and *Pseudomonas* spp. These genera can resist up to 90 to 140 Kg^-1^ of Cd showing their potential to use in soil bioremediation. In this study, multifunctional Cd-RB morphotypes were identified with the potential to develop integrative soil management of Cd-polluted soils.

## Supplementary Information

The online version contains supplementary material available at

## Acknowledgements

The authors thank the cacao farmers for letting them to collect samples. The authors thank the funding organization and the Universidad Nacional de Colombia for supporting the research group.

## Authors’ contributions

ETR, HCN, and JCZ conceived the project and designed the experiments. ETR, HCN and JCZ collected the samples. HCN and JCZ performed the experiments and analyzed the data. ETR supervised the research and analyzed the data. All authors wrote, and read and approved the final manuscript.

## Data availability

The datasets used and/or analyzed during the current study are available from the corresponding author on reasonable request. The 16S rRNA sequencing are available at NCBI according to GenBank accession numbers.

## Declarations

### Ethics approval and consent to participate

This work was conducted under Ministerio de Ambiente, Vivienda y Desarrollo Territorial (MAVDT) collection permit 0255, March 14^th^, 2014. Not applicable for animal or human data or tissue.

### Consent for publication

Not applicable.

