## Supplementary material for "Assessment of native cadmium-resistant bacteria in cacao (*Theobroma cacao* L.) - cultivated soils"

**Table 1.** Soil analysis of studied location in Nilo and Yacopí.

| Mun. | Local | pH | OC | Ca | K | Mg | Na | Al | ECEC | P | S | Cu | Fe | Mn | Zn | B | Cl | Sl | Sn | Text. |
|---|---|---|---|---|---|---|---|---|---|---|---|---|---|---|---|---|---|---|---|---|
| | | | % | Meq/100g | | | | | | Mg/kg | | | | | | | % | | | |
| Nilo | NC | 5.87 | 3.72 | 9.13 | 0.7 | 2.03 | 0.07 | 0.0 | 11.94 | >116 | 13.57 | 1.89 | 133.21 | 24.25 | 8.78 | <0.12 | 20 | 19 | 61 | Sandy Clay Loam |
| Yacopí | Y1 | 4.18 | 4.99 | 1.96 | 0.36 | 0.88 | 0.42 | 8.51 | 12.13 | 4.14 | 17.36 | 1.29 | 580.60 | 0.70 | 1.78 | 0.22 | 50 | 33 | 17 | Clay |
| | Y2 | 6.25 | 6.94 | 13.13 | 0.22 | 0.82 | 0.21 | 0.0 | 14.39 | 109.84 | 17.75 | 20.07 | 374.08 | 17.45 | 15.31 | 0.20 | 46 | 25 | 29 | Clay |
| | Y3 | 6.11 | 6.98 | 28.2 | 0.31 | 2.19 | 0.04 | 0.00 | 30.7 | 108 | 13.9 | 4.08 | 244 | 31.7 | 78.9 | 0.42 | 36 | 30 | 34 | Clay Loam |

Analyses were carried out by Soil Laboratory, Agricultural Sciences Faculty- Universidad Nacional de Colombia, Bogotá.

| PARAMETER | METHOD OF ANALYSIS | ASSESSMENT |
|---|---|---|
| **pH** | Soil suspension (1:1 w/v) | Potentiometric |
| **OC:** Organic carbon | Walkley-Black | Color |
| **N:** Total nitrogen | Based on OC (factor: 0.0862) | |
| **Ca, K, Mg:** Exchangeable bases | NH4 - acetate 1M pH 7 Extraction | Atomic absorption |
| **P:** Available phosphorous | Bray II | Color |
| **S:** Available sulfur | Extraction with monocalcium phosphate | Turbidity |
| **Cu, Fe, Mn, Zn:** Microelements | DTPA extraction | Atomic absorption |
| **B:** Boron | Extraction with monocalcium phosphate | Color |
| Clay (**Cl**), Silt (**Sl**), Sand (**Sn**) | Bouyoucos, dispersion with Na-Hexametaphosphate | Density |
| **Texture** | USDA triangular textural classification chart | |

**Table 2.** Morphological characterization at macro and microscopic levels of each isolated Cd-RB on culture media with 6 mg $kg^{-1}$ of Cd.

| Id[1] | | LOCAL. | COLOR (Oac) | | SHAPE | EDGE | SURFACE | ELEVATION | CONSISTENCY | GRAM STAINING | | ENDOSPORES | |
|---|---|---|---|---|---|---|---|---|---|---|---|---|---|
| | | | Obv. | Rev. | | | | | | Gram | Shape | Shape | Position |
| 1 | **NB2** | NC | 6 | 7 | Circular | Whole | Flat | Convex | Waxy | - | Bacillus | ND | |
| 2 | **NB10** | NC | 14 | 7 | Irregular | Wavy | Rough | Flat | Waxy | - | Bacillus | ND | |
| 3 | **NB40** | NC | 900 | 900 | Circular | Whole | Flat | Convex | Waxy | - | Bacillus | ND | |
| 4 | **NB58** | NC | 816 | 899 | Irregular | Wavy | Flat | Convex | Waxy | - | Bacillus | ND | |
| 5 | **NB59** | NC | 14 | 900 | Punctate | Whole | Flat | Convex | Waxy | - | Bacillus | ND | |
| 6 | **NB61** | NC | 857 | 900 | Irregular | Wavy | Rough | Convex | Waxy | - | Bacillus | ND | |
| 7 | **NB80** | NC | 857 | 900 | Punctate | Whole | Flat | Convex | Waxy | - | Bacillus | ND | |
| 8 | **YB11** | Y1 | 7 | 899 | Punctate | Whole | Flat | Convex | Waxy | - | Bacillus | ND | |
| 9 | **YB30** | Y1 | 7 | 900 | Punctate | Whole | Flat | Convex | Waxy | - | Bacillus | ND | |
| 10 | **YB42** | Y1 | 7 | 900 | Circular | Whole | Flat | Convex | Waxy | - | Bacillus | ND | |
| 11 | **YB43** | Y1 | 7 | 857 | Punctate | Whole | Flat | Convex | Waxy | - | Bacillus | ND | |
| 12 | **YB5** | Y2 | 5 | 30 | Irregular | Wavy | Flat | Flat | Waxy | - | Bacillus | ND | |
| 13 | **YB13** | Y2 | 857 | 7 | Punctate | Whole | Flat | Convex | Waxy | - | Bacillus | ND | |
| 14 | **YB22** | Y2 | 6 | 5 | Irregular | Wavy | Flat | Flat | Waxy | + | Bacillus | Circular | Term. |
| 15 | **YB48** | Y2 | 7 | 900 | Punctate | Whole | Flat | Convex | Waxy | - | Bacillus | ND | |
| 16 | **GB13** | Y3 | 909 | 900 | Circular | Whole | Flat | Convex | Waxy | - | Bacillus | ND | |
| 17 | **GB16** | Y3 | 909 | 909 | Circular | Whole | Flat | Convex | Waxy | - | Bacillus | ND | |
| 18 | **GB17** | Y3 | 899 | 7 | Punctate | Whole | Flat | Convex | Waxy | + | Bacillus | Circular | Term. |
| 19 | **GB18** | Y3 | 909 | 909 | Punctate | Whole | Flat | Convex | Waxy | + | Bacillus | Oval | Cent. |
| 20 | **GB58** | Y3 | 855 | 857 | Circular | Whole | Flat | Convex | Waxy | - | Bacillus | ND | |
| 21 | **GB66** | Y3 | 857 | 857 | Punctate | Whole | Flat | Convex | Waxy | - | Bacillus | ND | |
| 22 | **GB67** | Y3 | 855 | 847 | Punctate | Whole | Flat | Convex | Waxy | - | Bacillus | ND | |
| 23 | **GB68** | Y3 | 857 | 898 | Circular | Whole | Flat | Convex | Waxy | - | Bacillus | ND | |
| 24 | **GB71** | Y3 | 897 | 856 | Irregular | Wavy | Flat | Convex | Waxy | - | Bacillus | ND | |
| 25 | **GB73** | Y3 | 900 | 900 | Circular | Whole | Flat | Convex | Waxy | - | Bacillus | ND | |
| 26 | **GB78** | Y3 | 898 | 857 | Circular | Whole | Flat | Convex | Waxy | - | Bacillus | ND | |
| 27 | **GB82** | Y3 | 900 | 900 | Punctate | Whole | Flat | Convex | Waxy | - | Bacillus | ND | |
| 28 | **GB86** | Y3 | 898 | 899 | Circular | Whole | Flat | Convex | Waxy | - | Bacillus | ND | |
| 29 | **GB88** | Y3 | 899 | 900 | Circular | Whole | Flat | Convex | Waxy | - | Bacillus | ND | |
| 30 | **GB90** | Y3 | 859 | 900 | Circular | Whole | Flat | Convex | Waxy | - | Bacillus | ND | |

[1]ID: Morphotype identification; [2]Obverse (Obv.) and Reverse (Rev.) color of the petri dish according to Pantone scale (Oac); Local: Localization; ND: Not Determined; Term: terminal and Cent: Central.

**Table 3.** Molecular identification of isolated morphotypes in medium with 6 mg $kg^{-1}$ of Cd. Accession numbers represent the best match at the BLAST search.

| ID | Local. | Genus/Specie | Total Score | Query cover (%) | Ident (%) | E-value | Accession | Phylum | Order | Family |
|---|---|---|---|---|---|---|---|---|---|---|
| **NB2** | **NC** | *Pseudomonas aeruginosa* | 2512 | 100 | 99 | 0 | LT963552.1 | Proteobacteria | Pseudomonadales | Pseudomonadaceae |
| **NB10** | **NC** | *Burkholderia* | 2440 | 99 | 99 | 0 | KC756419.1 | Proteobacteria | Burkholderiales | Burkholderiaceae |
| **NB40** | **NC** | *Halomonas* | 1368 | 77 | 98 | 0 | MG386656.1 | Proteobacteria | Oceanospirillales | Halomonadaceae |
| **NB58** | **NC** | *Enterobacter* | 2431 | 100 | 99 | 0 | KR189808.1 | Proteobacteria | Enterobacteriales | Enterobacteriaceae |
| **NB59** | **NC** | *Serratia* | 17296 | 100 | 99 | 0 | LT907843.1 | Proteobacteria | Enterobacteriales | Enterobacteriaceae |
| **NB61** | **NC** | *Serratia* | 2468 | 100 | 99 | 0 | MH703475.1 | Proteobacteria | Enterobacteriales | Enterobacteriaceae |
| **NB80** | **NC** | *Serratia* | 2538 | 99 | 99 | 0 | EF550163.1 | Proteobacteria | Enterobacteriales | Enterobacteriaceae |
| **YB11** | **Y1** | *Enterobacter* | 2141 | 100 | 99 | 0 | MF666745.1 | Proteobacteria | Enterobacteriales | Enterobacteriaceae |
| **YB30** | **Y1** | *Serratia marcescens* | 17159 | 100 | 99 | 0 | CP027300.1 | Proteobacteria | Enterobacteriales | Enterobacteriaceae |
| **YB42** | **Y1** | *Serratia marcescens* | 17440 | 99 | 99 | 0 | CP018927.1 | Proteobacteria | Enterobacteriales | Enterobacteriaceae |
| **YB43** | **Y1** | *Serratia* | 1328 | 100 | 99 | 0 | MH703499.1 | Proteobacteria | Enterobacteriales | Enterobacteriaceae |
| **YB5** | **Y2** | *Enterobacter* | 18621 | 100 | 98 | 0 | CP012999.1 | Proteobacteria | Enterobacteriales | Enterobacteriaceae |
| **YB13** | **Y2** | *Pseudomonas* | 2494 | 100 | 99 | 0 | MH703511.1 | Proteobacteria | Pseudomonadales | Pseudomonadaceae |
| **YB22** | **Y2** | *Bacillus flexus* | 2518 | 99 | 98 | 0 | MH542283.1 | Firmicutes | Bacillales | Bacillaceae |
| **YB48** | **Y2** | *Enterobacter* | 2529 | 100 | 99 | 0 | MF101702.1 | Proteobacteria | Enterobacteriales | Enterobacteriaceae |
| **GB13** | **Y3** | *Burkholderia* | 2521 | 100 | 100 | 0 | MF795056.1 | Proteobacteria | Burkholderiales | Burkholderiaceae |
| **GB16** | **Y3** | *Burkholderia* | 2516 | 100 | 100 | 0 | FJ793552.1 | Proteobacteria | Burkholderiales | Burkholderiaceae |
| **GB17** | **Y3** | *Rhodococcus* | 1447 | 100 | 99 | 0 | KY697119.1 | Actinobacteria | Actinomycetales | Nocardiaceae |
| **GB18** | **Y3** | *Bacillus* | 1561 | 100 | 99 | 0 | KY622533.1 | Firmicutes | Bacillales | Bacillaceae |
| **GB58** | **Y3** | *Burkholderia* | 1323 | 100 | 99 | 0 | CP013429.1 | Proteobacteria | Burkholderiales | Burkholderiaceae |
| **GB66** | **Y3** | *Burkholderia* | 2536 | 100 | 99 | 0 | MT565307.1 | Proteobacteria | Burkholderiales | Burkholderiaceae |
| **GB67** | **Y3** | *Burkholderia* | 2436 | 100 | 99 | 0 | JF701980.1 | Proteobacteria | Burkholderiales | Burkholderiaceae |
| **GB68** | **Y3** | *Burkholderia* | 2516 | 100 | 99 | 0 | KY072913.1 | Proteobacteria | Burkholderiales | Burkholderiaceae |
| **GB71** | **Y3** | *Pseudomonas* | 1450 | 99 | 99 | 0 | MH517510.1 | Proteobacteria | Pseudomonadales | Pseudomonadaceae |
| **GB73** | **Y3** | *Pseudomonas* | 2569 | 100 | 99 | 0 | HG805731.1 | Proteobacteria | Pseudomonadales | Pseudomonadaceae |
| **GB78** | **Y3** | *Pseudomonas putida* | 2538 | 100 | 100 | 0 | MH084903.1 | Proteobacteria | Pseudomonadales | Pseudomonadaceae |
| **GB82** | **Y3** | *Burkholderia* | 2512 | 100 | 100 | 0 | CP016001.1 | Proteobacteria | Burkholderiales | Burkholderiaceae |
| **GB86** | **Y3** | *Pseudomonas* | 2412 | 100 | 99 | 0 | JQ951925.1 | Proteobacteria | Pseudomonadales | Pseudomonadaceae |
| **GB88** | **Y3** | *Pseudomonas* | 2519 | 100 | 99 | 0 | KY460976.1 | Proteobacteria | Pseudomonadales | Pseudomonadaceae |
| **GB90** | **Y3** | *Herbaspirillum* | 1192 | 99 | 99 | 0 | FJ687986.1 | Proteobacteria | Burkholderiales | Oxalobacteraceae |

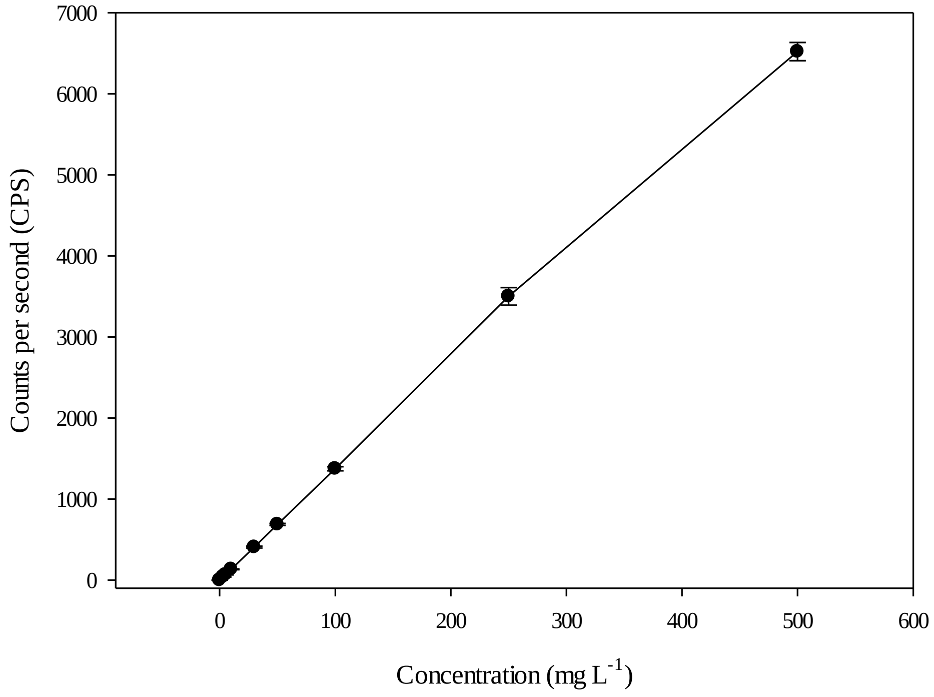


**Fig. 1.** Calibration curve for Cd determination using eight concentrations and three lectures per concentration at different times.

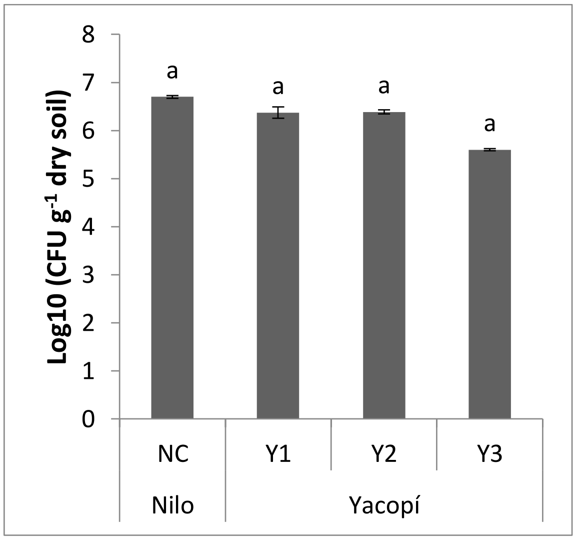


**Fig. 2.** Log10CFU $g^{-1}$ dry soil in culture media with 6 mg $Kg^{-1}$ of Cd, isolated from Nilo and Yacopí soils. Letters over the bars represent significant differences between locations ($p<0.05$) according to Tukey's test.
